# RBM15::MKL1 uncouples SPOC-dependent m^6^A remodeling from WTAP recruitment

**DOI:** 10.64898/2026.11.13.766277

**Authors:** Jether Amos Espinosa, Mi Chen, Madeline Y. Mayday, Hummi Mohammed, Gina Lee, Lin Wang, Ping-Xia Zhang, Nadia Ayala-Lopez, Diane S. Krause

## Abstract

The recurrent t(1;22) translocation associated with pediatric acute megakaryoblastic leukemia generates the RBM15::MKL1 (RM) fusion protein that retains the RNA-binding and SPOC domains of RBM15. RBM15 recruits the *N*^6^-methyladenosine (m^6^A) methyltransferase complex (MTC), of which WTAP is a key regulatory component. How RM engages this machinery and whether the SPOC domain is required for its leukemogenic activity remain unclear. Here, we investigated the molecular determinants of WTAP recruitment by RBM15 and RM and their contribution to RM-mediated disruption of megakaryocyte differentiation. Structure-guided mutagenesis identified a conserved basic surface within the isolated RBM15 SPOC domain that is required for WTAP association. Unexpectedly, disruption of this surface or complete SPOC deletion failed to abolish WTAP association with full-length RBM15 or RM, indicating that additional determinants mediate MTC recruitment. In human erythroleukemia cells, RM impaired TPA-induced megakaryocytic maturation, as assessed by differentiation markers, polyploidization, and morphology; this blockade persisted following SPOC disruption. However, transcriptomic analysis demonstrated that RM lacking the SPOC domain elicited an attenuated gene expression response compared with unmutated RM, including alterations in WNT/β-catenin-associated genes. m^6^A-enriched RNA profiling further revealed extensive remodeling of the m^6^A epitranscriptome by RM, while SPOC deletion shifted RM-associated m^6^A changes toward a more wild-type-like state. Together, these findings demonstrate that although the SPOC domain contributes to RM-mediated molecular regulation, it is dispensable for stable MTC association and maintenance of the differentiation blockade, revealing previously unrecognized redundancy in RBM15-mediated recruitment of the m^6^A machinery.

**Data Sharing Statement:** All data generated in this study are included in this article and its supplementary information. RNA-seq and EpiPlex data have been deposited in the Gene Expression Omnibus (GEO) repository and are accessible with the accession number GSE349759.

**Key Points:**

- SPOC deletion shifts RM-associated m^6^A changes and gene expression toward a wild-type-like state without restoring megakaryocyte maturation.
- The RBM15 SPOC domain is dispensable for WTAP recruitment by full-length RBM15 and the leukemogenic RBM15::MKL1 (RM) fusion.

## Introduction

Acute megakaryoblastic leukemia (AMKL) is a rare but aggressive subtype of acute myeloid leukemia (AML) characterized by the malignant expansion of immature megakaryoblasts.^1,2^ AMKL accounts for approximately 10% of pediatric AML cases and encompasses biologically distinct subtypes characterized by recurrent chromosomal rearrangements and fusion oncogenes.^3,4^ Among these, the RBM15::MKL1 (RM) fusion generated by the t(1;22)(p13;q13) translocation retains all functional domains of endogenous RBM15 and MKL1 (**Figure 1A**) and, as a hallmark of AMKL,^5,6^ is associated with impaired megakaryocytic differentiation.^7–9^ The lack of consensus on optimal AMKL treatment with intensive chemotherapy versus hematopoietic stem cell transplantation^10^ underscores the need to define the molecular mechanisms that sustain leukemogenesis and identify unique therapeutic vulnerabilities.

**Figure 1.**
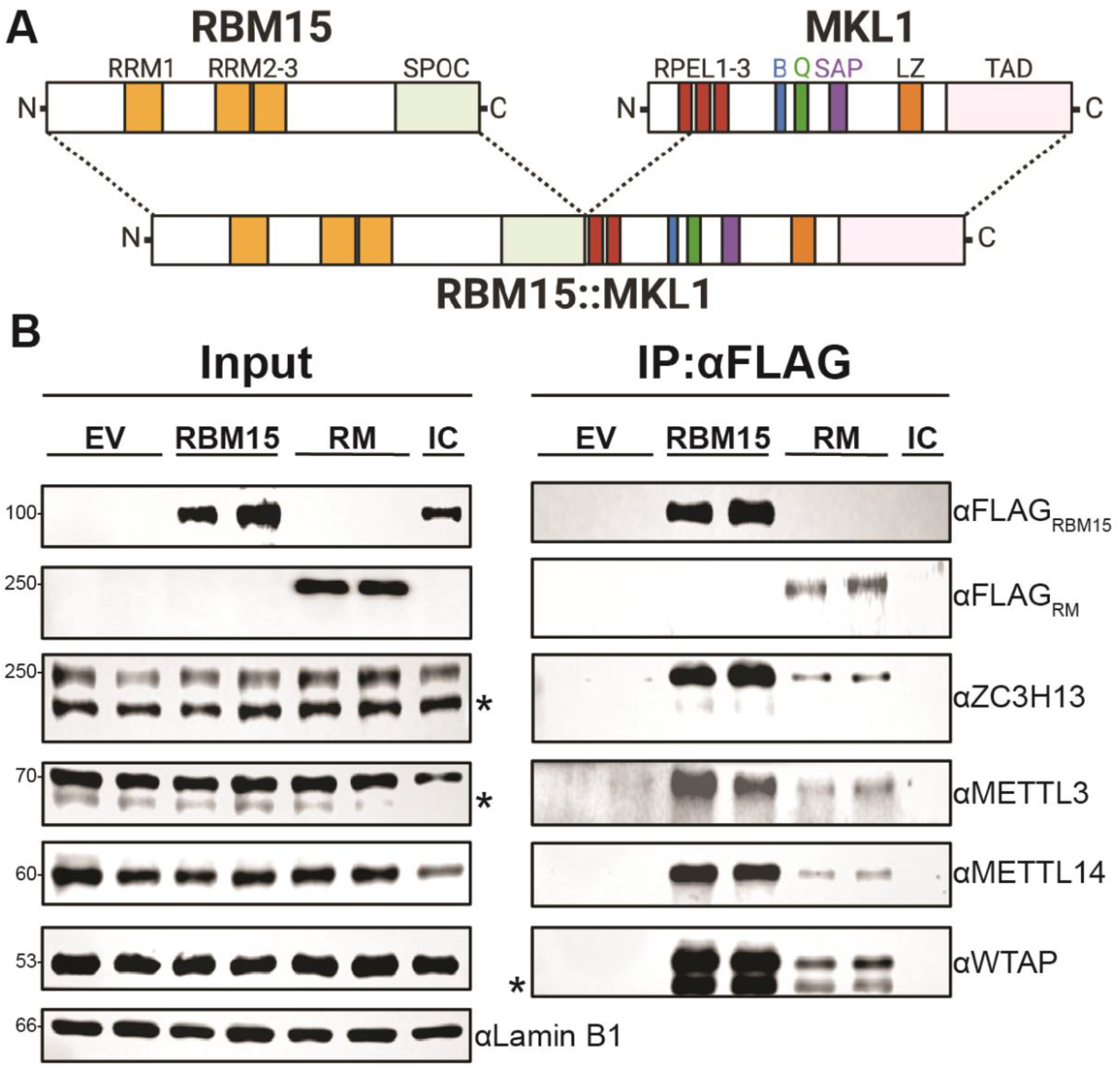
RBM15::MKL1 (RM) retains association with the m^6^A methyltransferase complex. (**A**) Schematic representation of the domain architecture of RBM15, MKL1, and the RM fusion protein. RM retains the three RNA-recognition motifs (RRM1-3) and Spen paralog and ortholog C-terminal (SPOC) domain of RBM15, together with RPEL2-3 motifs, basic region (B), glutamine-rich region (Q), SAP domain, leucine zipper (LZ), and transactivation domain (TAD) of MKL1. Image created using BioRender.com. (**B**) Co-immunoprecipitation (co-IP) using anti-FLAG pulldown of either FLAG-tagged RBM15 or RM compared to empty vector (EV) control followed by immunoblotting to detect members of the m^6^A methyltransferase complex (i.e., ZC3H13, METTL3, METTL14, and WTAP) in doxycycline (dox)-induced HEL cells. Co-IP was performed from nuclear lysates with Lamin B1 serving as a loading control. Representative immunoblots from three independent biological replicates are shown (n = 3). Asterisks (*) indicate non-specific bands. IgG isotype control (IC) for the pulldown is from nuclear lysate of dox-induced HEL-RBM15.

RM promotes leukemic transformation by enhancing the expression of *Frizzled* transcripts that support leukemic self-renewal.^8^ Dysregulation of *N*^6^-methyladenosine (m^6^A), the most abundant epitranscriptomic modification on mRNA^11,12^, is also implicated in RM-driven leukemogenesis.^8^ Studies using the murine 6133 AMKL cell line, derived from a knock-in model expressing murine *Rbm15* fused to human *MKL1*, demonstrated that continued RM expression is required for leukemic viability and proliferation and is associated with increased m^6^A modification of select RNA targets associated with leukemogenic transcriptional programs.^8^ Consistent with this model, pharmacologic inhibition of the m^6^A methyltransferase complex (MTC) suppresses leukemia progression in preclinical models and promotes megakaryocytic differentiation, highlighting the m^6^A pathway as a potential therapeutic target.^8,13^

RBM15, the N-terminal component of RM, can act as a link between RNA and the MTC.^14,15^ Its N-terminal RNA recognition motifs (RRMs) bind specific mRNA transcripts^16,17^, while its Spen paralog and ortholog C-terminal (SPOC) domain associates with the MTC regulatory subunit WTAP^18,19^ and facilitates the localization of the catalytic METTL3-METTL14 heterodimer to target mRNAs.^20,21^ Structural and biochemical studies identified a conserved positively charged surface within the RBM15 SPOC domain that mediates phosphoserine-dependent protein-protein interactions and is thought to affect WTAP binding.^19^ The prevailing model therefore proposes that SPOC-dependent WTAP recruitment supports RBM15-directed m^6^A deposition, and by extension, RM leukemogenic activity.

Whether SPOC is required for WTAP association with full-length RBM15 or RM, or for RM-mediated inhibition of megakaryocytic differentiation, remains unclear. Here, we used structure-guided mutagenesis and biochemical studies to investigate SPOC-dependent WTAP association, m^6^A regulation, and megakaryocytic differentiation in a model of RM-AMKL.

## Methods

### Cell culture and morphology assay

Human embryonic kidney 293T (HEK293T; ATCC, Manassas, VA) cells were cultured in DMEM (Thermo Fisher Scientific, Waltham, MA) and human erythroleukemia (HEL; ATCC) cells in RPMI-1640 (Thermo) medium, each supplemented with 10% (v/v) fetal bovine serum (FBS; Takara Bio, San Jose, CA), 1% (v/v) GlutaMAX (Thermo), 100 U/mL penicillin, and 100 µg/mL streptomycin (Thermo). Tetracycline-free version of FBS (Takara) were used in culturing HEL cells. Cells were maintained at 37°C in a humidified atmosphere containing 5% CO_2_. Cell morphology was assessed using cytospins stained using a modified Romanowsky Differential Quik III Stain (Polysciences, Warrington, PA). Images were acquired by brightfield microscopy at 20× magnification using the Olympus BX51 Microscope (Evident Scientific, Waltham, MA).

### Co-immunoprecipitation (co-IP) and immunoblotting

Nuclear lysates were prepared using the NE-PER Nuclear and Cytoplasmic Extraction Reagents (Thermo) as described by the manufacturer. For co-IP, lysates were incubated with FLAG-conjugated Dynabeads Protein G (Thermo) overnight at 4°C, washed, and eluted. Proteins were separated by SDS-PAGE (Bio-Rad Laboratories, Hercules, CA), transferred to nitrocellulose (Bio-Rad), and detected using primary and secondary horseradish peroxidase-conjugated antibodies incubated at RT for 1 hr, and visualized using the G:Box Chemi-XRQ (Syngene, Cambridge, United Kingdom). Antibodies and dilutions are listed in **Supplementary Table S1**.

### RNA isolation

Total RNA was isolated using RNeasy Kit (Qiagen, Venlo, The Netherlands) with on-column DNase digestion (Qiagen), and 5.0 µg of total RNA was subjected to poly(A) enrichment using the NEBNext poly(A) mRNA Magnetic Isolation Module (New England Biolabs, Ipswich, MA) as described by the manufacturer.

### EpiPlex RNA assay and EpiScout analysis

Epitranscriptomic profiling was conducted using the EpiPlex RNA Mod Encoding Kit (AlidaBio, San Diego, CA) using 50.0 ng of poly(A)-enriched and DNase I-treated RNA as input. Prior to m^6^A enrichment onto EpiPlex beads, the purified mRNA was mixed with spike-in controls, fragmented, end-repaired, and ligated to an Illumina P7 adapter. Approximately 10% of the prepared RNA was allocated to non-enriched solution control libraries, while the remaining 90% was incubated with EpiPlex beads containing m^6^A-specific RNA binders for enrichment. At the bead surfaces, captured RNA fragments were randomly primed by proximally anchored adapters containing Illumina P5 adapters, a unique molecular identifier (UMI), and a modification barcode (MBC) which was replicated into cDNA by reverse transcriptase. Sample indices were added to each sample using PCR via the EpiPlex Unique Dual Index Primer Kit (AlidaBio). Libraries were sized on the Agilent TapeStation (Agilent Technologies, Santa Clara, CA) and quantified by Qubit (Thermo). Separate library pools containing equal representation of each library were made for bead-enriched and solution control libraries. Each library was sequenced to a target depth of ∼30 million 100-bp paired-end reads on an Illumina NovaSeq X Plus (Illumina, San Diego, CA). Sequencing data were analyzed using the EpiScout RNA Analysis Suite (AlidaBio), with reads aligned to the human reference genome GRCh38.p14 according to the analysis pipeline available on the manufacturer’s website. EpiScout outputs were used to assess sequencing and modification-specific quality-control metrics (**Figure S5A-D**). Differential and intersecting m^6^A modification regions generated by EpiScout were used for comparisons between experimental conditions.

### Flow cytometry

Megakaryocytic differentiation was assessed using anti-CD49b conjugated to either APC or PE for cell surface markers (BioLegend, San Diego, CA) and propidium iodide (PI; BioLegend) for DNA content to assess for polyploidization. Cells were stained with anti-CD49b antibody at 4°C for 30 min protected from light. After staining, cells were washed with FACS buffer and fixed using the BD Phosflow Lyse/Fix Buffer (Waters Biosciences, Milford, MA) as described by the manufacturer, followed by digestion with 20 ng/mL RNase A (Thermo) at 37°C, 5% CO_2_ in a humidified incubator, and FACS tubes containing cell suspensions were agitated by flicking every 30 min for 3 h total. Samples were analyzed on an LSRII or LSRFortessa flow cytometer (Waters) and processed in FlowJo (Waters). Antibodies and dilutions are listed in **Supplementary Table S1**.

### Stable transfection of HEL cells

To establish HEL cells with inducible RM (HEL-RM), parental HEL cells were transfected, as described.^9^ Single-colony clones grown in semisolid medium with high expression of RM were isolated and expanded. All HEL-RM mutant variants were generated using the same methods.

### Transient transfection of HEK293T cells

Cells were transiently transfected with FuGENE HD (Promega, Fitchburg, WI) as described by the manufacturer and cultured for 72 h. Expression of recombinant proteins was confirmed by immunoblotting.

### Statistical analysis

Performed using GraphPad Prism version 11.0.2 (GraphPad Software, San Diego, CA). Data are presented as mean ± standard deviation or standard error of the mean as indicated. Statistical tests are specified in figure legends; \**P*-values < 0.05 were considered statistically significant (\*\**P* < 0.01; \*\*\**P* < 0.001; \*\*\*\**P* < 0.0001). n = not significant.

### Structural modeling of full-length RBM15 and RBM15 SPOC domain

Structures of full-length RBM15, truncated RBM15, and isolated RBM15 SPOC domain were predicted using AlphaFold3 (https://alphafoldserver.com/welcome) and visualized using UCSF ChimeraX (https://www.cgl.ucsf.edu/chimerax/).^22^

## Results

### RM retains association with the m^6^A MTC

RM retains the three N-terminal RRMs and the C-terminal SPOC domain of RBM15 together with the domains contributed by MKL1 (**Figure 1A**). RBM15 functions as an RNA-binding adaptor for the m^6^A writer machinery and interacts with WTAP, which associates with the METTL3-METTL14 catalytic core and other MTC regulators.^15,20,21,23–25^ To determine whether RM preserves these interactions, we examined MTC association with full-length RBM15 and RM in dox-inducible HEL cells. Anti-FLAG immunoprecipitation (IP) showed that both RBM15 and RM associate with endogenous WTAP, METTL3, METTL14, and ZC3H13 (**Figure 1B, S1A**). In contrast, MKL1 did not detectably associate with WTAP in dox-inducible HEL-MKL1 cells (**Figure S1B**), indicating that WTAP association is not conferred by the MKL1 portion of the fusion. Together, these findings demonstrate that fusion to MKL1 does not preclude MTC association.

### The isolated RBM15 SPOC domain contains a conserved WTAP-binding interface

Because the RBM15 SPOC domain is retained in RM and can bind directly to WTAP,^19^ we next defined this interaction interface and tested its requirement in full-length RBM15 and RM. FLAG-tagged RBM15 truncations encompassing the N-terminal RRMs, intrinsically disordered region (IDR), or the C-terminal SPOC domain (**Figure 2A**) were expressed in HEK293T cells and assessed for endogenous WTAP co-IP. Full-length RBM15 and the isolated SPOC-containing C-terminal fragments robustly associated with WTAP (**Figure 2B, S2D**). In contrast, whereas the N-terminal fragments spanning amino acid 1-507, which contain most of the RRM1-3 region but lack the SPOC domain, did not associate with WTAP. These findings localized WTAP-binding activity to the SPOC-containing C-terminal region in the tested fragments. However, because the N-terminal constructs lacked the additional sequences beyond SPOC, these results did not establish that the SPOC domain was uniquely required for WTAP association in full-length RBM15.

**Figure 2.**
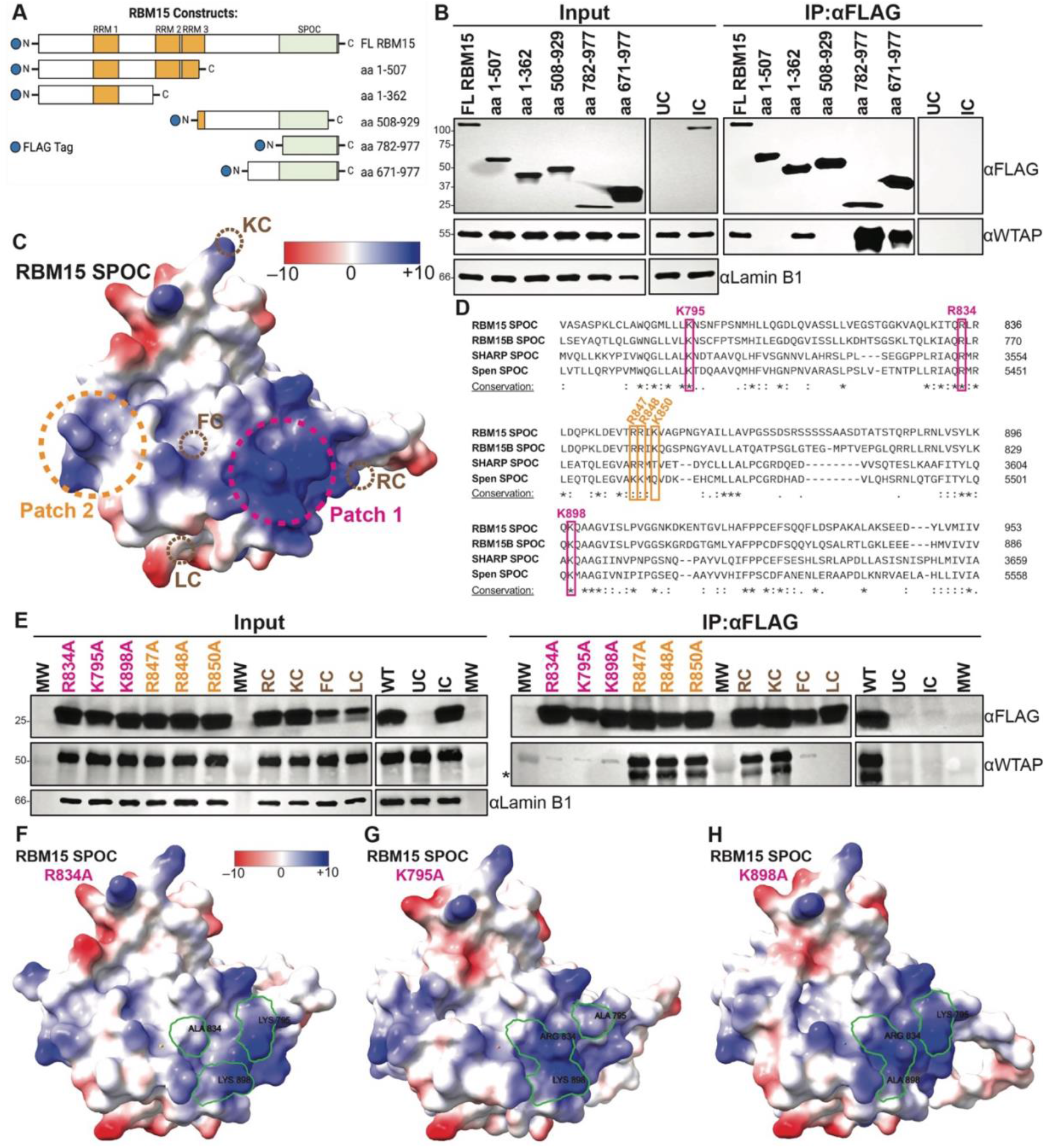
The isolated RBM15 SPOC domain contains a conserved WTAP-binding interface. (**A**) Cartoon depiction starting with full-length (FL) RBM15 and its three N-terminal RNA recognition motifs (RRM1, RRM2, and RRM3) in yellow, Spen paralog and ortholog C-terminal (SPOC) domain in green, and intrinsically disordered regions in white. Truncated constructs with corresponding amino acid (aa) positions. All constructs are N-terminally FLAG tagged (blue circle). Image created using BioRender.com (**B**) Co-immunoprecipitation (co-IP) using anti-FLAG pulldown of FL RBM15 and truncated constructs compared to untransfected control (UC) followed by immunoblotting to detect WTAP in HEK293T cells. Nuclear fraction of cell lysates was used for co-IP and thus normalized to Lamin B1 loading control. IgG Isotype control (IC) for the pulldown is from nuclear lysate of HEK293T transfected with FL RBM15 construct. Representative blot is shown (n = 3 biological replicates). (**C**) AlphaFold3 (AF3) structural model of the RBM15 SPOC domain. Electrostatic surface potential was visualized by using the Coulombic Surface Coloring tool in UCSF ChimeraX^22^ and is depicted ranging from –10 (red), 0 (white), to +10 (blue) kcal/(mol*e). The conserved basic Patch 1 is indicated by the dashed fuchsia circle (right) while the moderately conserved basic Patch 2 is indicated by the dashed orange circle (left). (**D**) Multiple sequence alignment via Clustal Omega^44^ of SPOC domains from human (RBM15, RBM15B, and SHARP) and *Drosophila* (Spen). Colored squares in fuchsia and orange correspond to basic residues in Patch 1 and Patch 2, respectively. Conservation is denoted as high (*), moderate (:), and low (.). (**E**) Co-IP using anti-FLAG pulldown of RBM15 SPOC domain alanine mutants on Patch 1 (fuchsia) and Patch 2 (orange) compared with unmutated SPOC (WT) and UC, followed by immunoblotting to detect WTAP in HEK293T cells. Non-Patch 1 alanine mutant controls (purple) include R886A (RC), K912A (KC), F923A (FC) and L933A (LC). Nuclear fraction of cell lysates was used. IC for the pulldown is from nuclear lysate of HEK293T transfected with WT construct. Molecular weight (MW) wells are indicated with protein mass in kDa. Representative blot is shown (n = 3 biological replicates). Asterisk (*) indicates non-specific bands. (**F**-**H**) AF3 structural model of individual Patch 1 mutations on the RBM15 SPOC domain visualized as in (C). Loss of Patch 1 electrostatic potential is outlined in green in all three mutants.

An AlphaFold 3 (AF3)^26^ model of RBM15 recapitulated the overall fold and surface features of the previously reported RBM15 SPOC crystal structure^19^, including two conserved positively charged pockets termed Patch 1 and Patch 2 (**Figure 2C-D**). Alanine scanning mutagenesis targeting each patch, one residue at a time, showed that individual Patch 1 alanine mutants markedly reduced WTAP association, whereas Patch 2 mutations had minimal effect, similar to control mutations outside either Patch 1 or 2 (**Figure 2E and S2A, B, E, and F**). K898A and K795A alanine mutants in Patch 1 of full-length Rbm15 abrogate binding to Setd1b^27^, a member of the histone methyltransferase complex that methylates H3K4, consistent with the SPOC domain binding other proteins in addition to WTAP.

AF3 electrostatic models showed that the prominent positive surface potential centered on Patch 1 was diminished by individual R834A, K795A, and K898A substitutions (**Figure 2F-H**) and further neutralized by the triple Patch 1 mutant (**Figure S2C**). Control substitutions outside Patch 1 did not measurably impair WTAP co-IP (**Figure S2A, B, and F**), indicating that loss of binding was specific to the Patch 1 interface.

Collectively, these findings confirm that RBM15 SPOC domain binds WTAP and identify the conserved Patch 1 basic surface as the canonical molecular interface that mediates this protein-protein interaction. Because these studies used the isolated SPOC domain, we next tested whether the same molecular requirements govern WTAP recruitment by full-length RBM15.

### Full-length RBM15 retains WTAP association despite disruption of the SPOC domain

We next tested whether Patch 1 is required for WTAP association in full-length RBM15. In contrast to isolated SPOC, full-length RBM15 carrying R834A, K795A, and K898A retained WTAP association comparable to wild-type RBM15 (**Figure 3A-B**). Simultaneous mutation of all three Patch 1 residues produced only a modest reduction, with substantial interaction remaining. Complete SPOC deletion (ΔSPOC) likewise did not completely abolish WTAP association, indicating that determinants outside SPOC contribute to WTAP recruitment by full-length RBM15.

**Figure 3.**
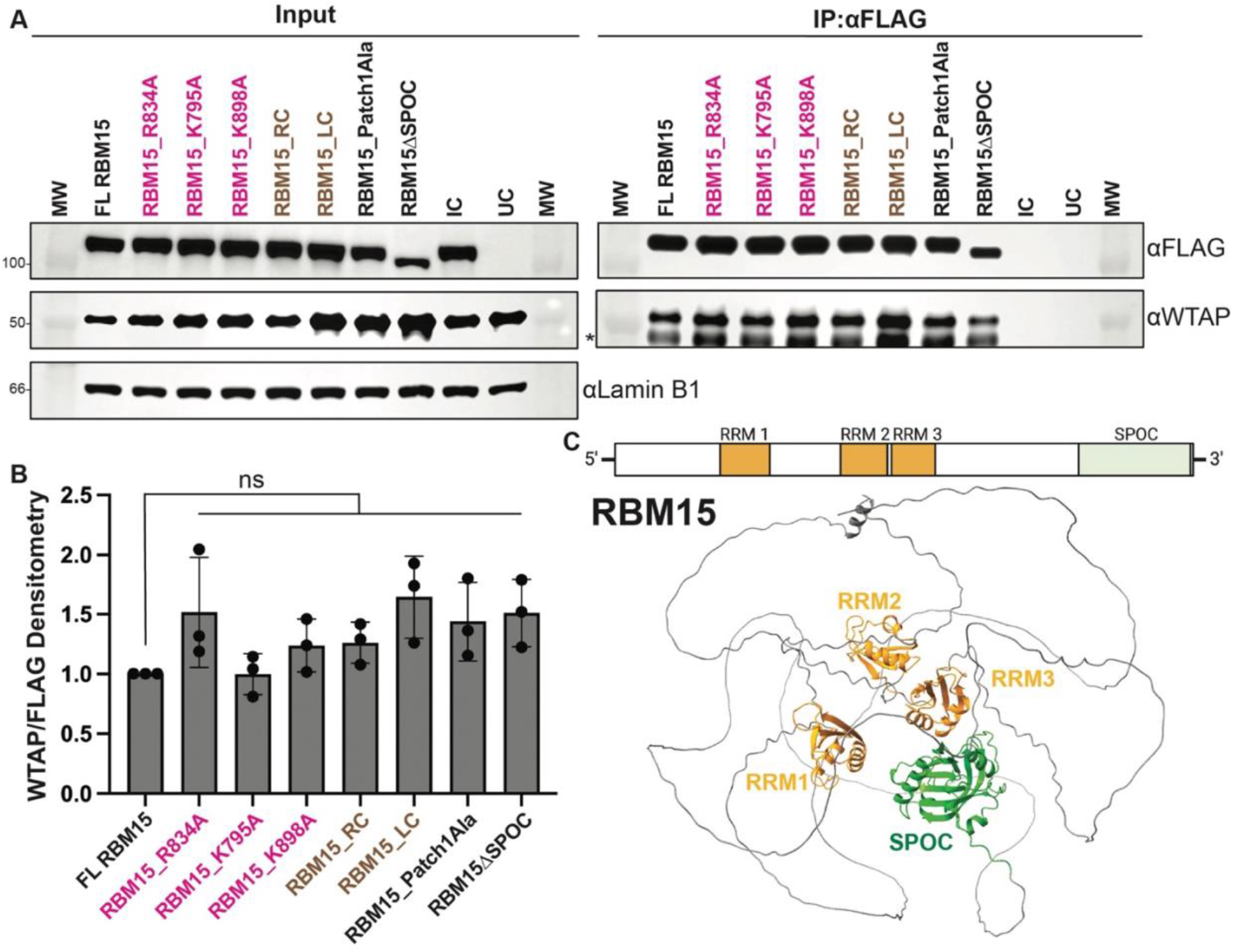
RBM15 retains WTAP association despite disruption of the SPOC domain. (**A**) Co-IP using anti-FLAG pulldown of full-length (FL) RBM15 and corresponding alanine mutants on Patch 1 (fuchsia), triple Patch 1 alanine mutant (Patch1Ala), and SPOC domain deletion (ΔSPOC) compared with an untransfected control (UC), followed by immunoblotting to detect WTAP in HEK293T cells. Nuclear fraction of cell lysates was used. IgG Isotype control (IC) for the pulldown is from nuclear lysate of HEK293T transfected with FL RBM15 construct. Non-Patch 1 alanine mutant controls on FL RBM15 (brown) include R886A (RC) and L933A (LC). Molecular weight (MW) wells are indicated with protein mass in kDa. Representative blot is shown (n = 3 biological replicates). Asterisk (*) indicates non-specific bands. (**B**) Densitometric quantification of WTAP co-IP from (A) normalized to FLAG pulldown and expressed relative to FL RBM15 (FL = 1.0). Data are presented as mean ± standard error of the mean (SEM). Statistical significance was assessed using ordinary one-way ANOVA with Dunnett’s multiple comparisons test versus WT (ns, not significant). (**C**) AlphaFold3 structural model of full-length RBM15 visualized as a ribbon model in UCSF ChimeraX highlighting three N-terminal RNA recognition motifs (RRM1, RRM2, and RRM3) in yellow and the Spen paralog and ortholog C-terminal (SPOC) domain in green.

To explore a structural basis for this finding, AF3 was used to predict the structure of full-length RBM15, which remains the only member of the MTC lacking an experimentally determined structure (though the SPOC domain structure has been determined).^21^ AF3 modeling positioned the N-terminal RRMs near the C-terminal SPOC domain, including a potential RRM1-SPOC interface (**Figure 3C**). RRM2 and RRM3 of RBM15 are known to adopt a coaxial arrangement that flanks RNA,^16^ and RRMs more broadly can also mediate protein-protein interactions.^28^ Thus, RRM1 or the two IDRs that flank it may contribute directly or indirectly to WTAP recruitment.

Together, these results support a revised model in which full-length RBM15 possesses additional mechanisms for recruiting WTAP beyond the SPOC-mediated interaction.

### RM similarly recruits WTAP independent of the SPOC domain

We next tested whether this finding is extended to RM. To address this question in a disease-relevant context, we generated dox-inducible HEL cell lines expressing FLAG-tagged wild-type RM or corresponding SPOC domain variants. HEL cells provide a well-established model of megakaryocytic differentiation undergoing characteristic maturation following treatment with 12-*O*-tetradecanoylphorbol-13-acetate (TPA),^29,30^ thereby enabling mechanistic studies of RM function within a physiologically relevant hematopoietic setting.

Using site-directed mutagenesis, the same structure-guided mutations previously examined in full-length RBM15 were introduced in RM, including individual alanine substitutions targeting conserved Patch 1 basic residues, a triple Patch 1 alanine mutant, and a complete SPOC domain deletion. Stable dox-inducible HEL cell lines were generated for each construct and comparable expression of FLAG-tagged RM proteins following dox induction was confirmed by immunoblotting prior to interaction studies (**Figure 4A**).

**Figure 4.**
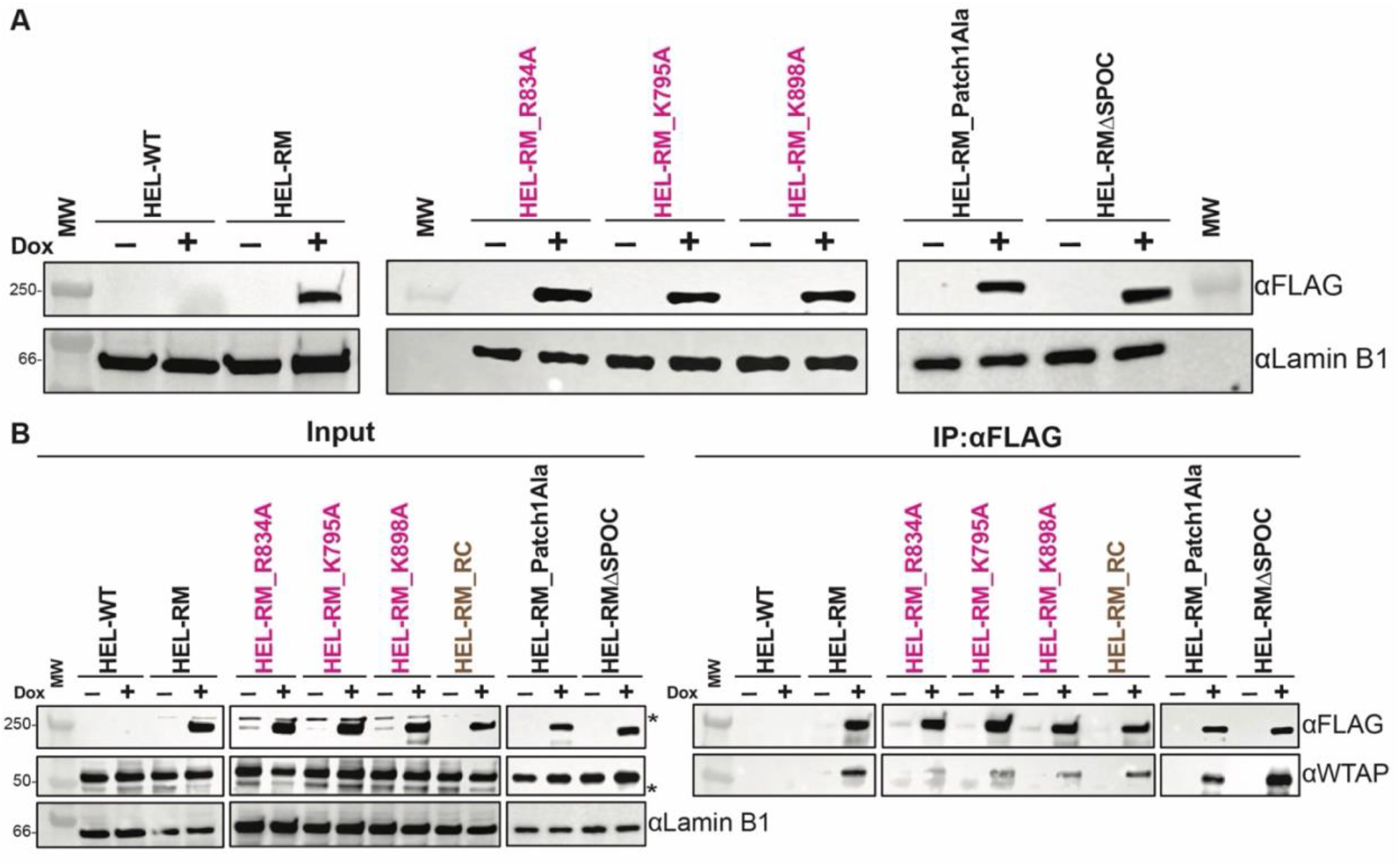
RM recruits WTAP independent of the SPOC domain. (**A**) Representative immunoblot confirming doxycycline (dox)-inducible expression of FLAG-tagged unmutated and mutant RM constructs in HEL cell lines. Mutated RM constructs include individual Patch 1 alanine mutants (fuchsia), the triple Patch 1 alanine mutant (Patch1Ala), and a SPOC domain-deleted mutant (ΔSPOC). Nuclear fractions from the indicated HEL cell lines cultured in the absence (–) or presence (+) of dox were probed with the indicated antibodies. Lamin B1 was used as the nuclear loading control. Molecular weight (MW) wells are indicated with protein mass in kDa. Representative blot is shown (n = 3 biological replicates). (**B**) Co-immunoprecipitation (co-IP) of dox-induced HEL cells from (A) using anti-FLAG pulldown followed by immunoblotting to detect WTAP. R886A (RC), a non-Patch 1 alanine mutant control on RM (brown), was included. Nuclear fraction of cell lysates was used for co-IP and thus normalized to nuclear Lamin B1 loading control. Representative blot is shown (n = 3 biological replicates). Asterisks (*) indicate non-specific bands.

To determine whether disruption of the SPOC domain altered recruitment of the MTC, FLAG-tagged proteins were immunoprecipitated from dox-induced nuclear HEL cell lysates and analyzed for endogenous WTAP association by co-IP. Consistent with our observation in full-length RBM15, none of the individual Patch 1 alanine substitutions impaired WTAP association within the context of the RM fusion protein (**Figure 4B, S3**). Likewise, neither simultaneous mutation of all three conserved Patch 1 residues, nor complete deletion of the SPOC domain abrogated WTAP co-IP (**Figure 4B, S3**).

These findings demonstrate that the fusion protein mirrors the behavior of full-length RBM15, retaining robust interaction with WTAP despite disruption of the previously characterized SPOC-dependent binding interface.^19^ Together with our preceding analyses, these data indicate that although the SPOC domain constitutes the principal WTAP-binding surface as an isolated domain, it is dispensable for stable WTAP association in the context of the full-length protein architecture. These observations establish that loss of the SPOC domain-WTAP interaction alone is unlikely to abolish m^6^A writer complex recruitment by the leukemogenic fusion protein, motivating subsequent experiments to determine whether disruption of this interface nevertheless alters RM-mediated inhibition of megakaryocytic differentiation, RNA expression and/or m^6^A changes.

### SPOC domain disruption fails to rescue megakaryocytic differentiation

To determine whether the disruption of the SPOC-WTAP interaction rescues the differentiation block imposed by RM, we evaluated megakaryocytic maturation using the same dox-inducible HEL cell model. Following induction of RM expression, cells were treated with TPA to promote megakaryocytic differentiation and were subsequently analyzed by complementary phenotypic, morphological, and biochemical approaches. Megakaryocytic differentiation was first assessed by flow cytometric analysis of cell surface CD49b, which is upregulated with megakaryocyte maturation.^29,31–33^ Consistent with published data^8^, TPA treatment induced robust CD49b expression in control cells, whereas expression of RM markedly suppressed this differentiation-associated increase (**Figure 5A**). SPOC disruption failed to reverse this phenotype. Specifically, HEL cells expressing the individual Patch 1 mutants (R834A, K795A, and K898A), the triple Patch 1 mutant, or the SPOC domain deletion all exhibited the same inhibition of TPA-induced CD49b expression as unmutated RM-expressing cells (**Figure 5B**). Flow cytometry demonstrated a persistent reduction in the CD49b-high population across all RM variants (compare blue versus red lines), indicating that the loss of SPOC-mediated WTAP interaction is insufficient to affect the block in megakaryocyte differentiation.

**Figure 5.**
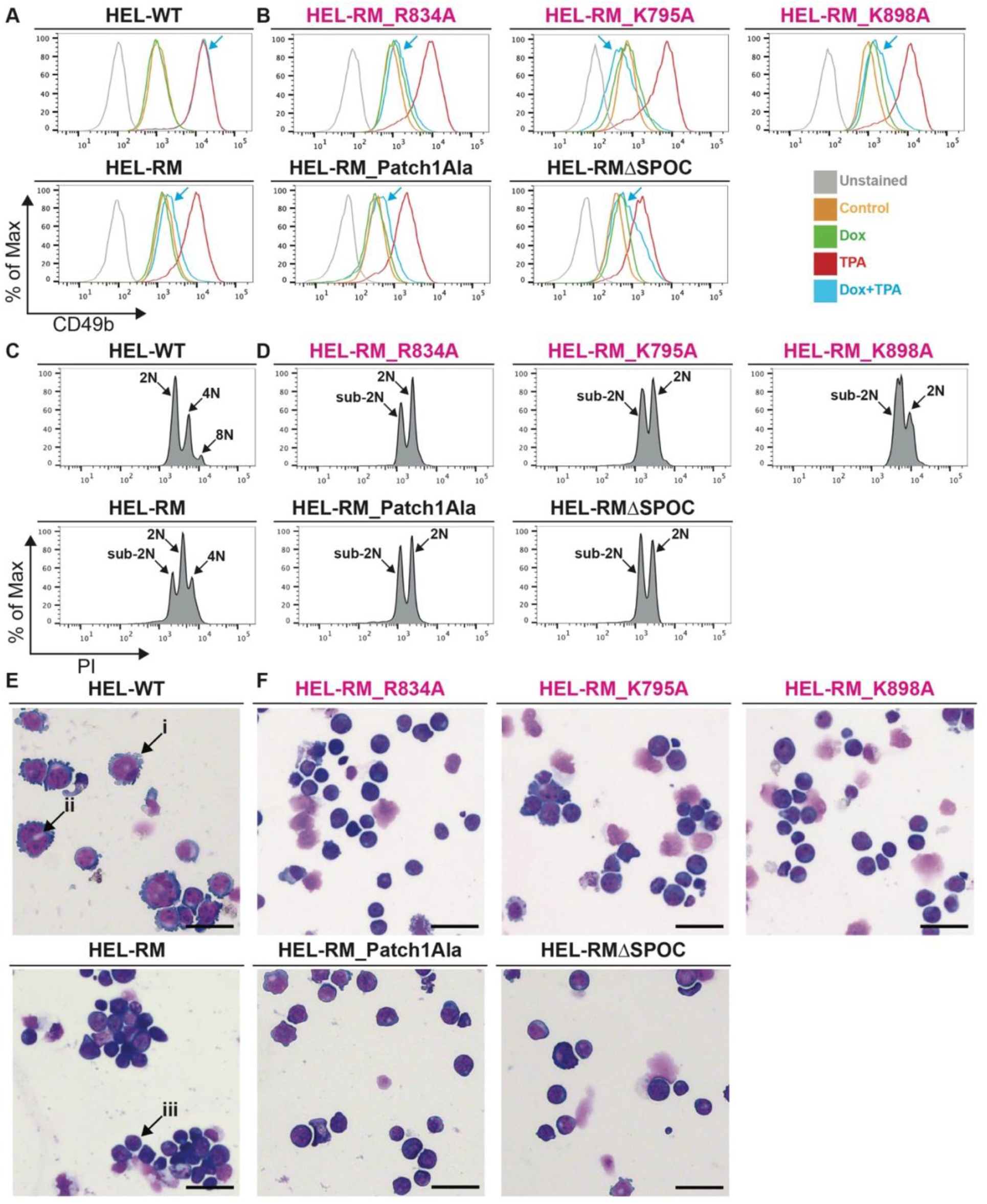
SPOC domain disruption fails to rescue megakaryocytic differentiation. (**A**) CD49b expression of HEL-WT (top) and HEL-RM (bottom) cells cultured without doxycycline (dox; orange), with dox (green), with 12-*O*-tetradecanoylphorbol-13-acetate (TPA; red), or with both dox and TPA (blue with arrow), relative to unstained control (grey). (**B**) CD49b expression of HEL cells with dox-induced individual Patch 1 RM mutants (top row, fuchsia), triple Patch 1 alanine mutant (Patch1Ala; bottom row), and SPOC domain-deleted mutant (ΔSPOC; bottom row) with the same conditions as (A). (**C**) Flow cytometry histograms showing ploidy peaks of propidium iodide (PI)-stained HEL-WT (top) and HEL-RM (bottom) cells treated with dox and TPA for 72 hours. Ploidy peaks of 2N, 4N, and 8N are noted in HEL-WT as reference. Presence of a sub-2N peak is noted in HEL-RM as reference. (**D**) Ploidy histograms for HEL-RM Patch 1 single alanine mutants (top row, fuchsia), triple Patch1 alanine mutant (Patch1Ala; bottom row), and SPOC domain-deleted mutant (ΔSPOC; bottom row). All cell lines were dox- and TPA-induced for 72 hours as in (C). (**E**) Representative Differential Quik III (modified Romanowsky)-stained cytospins of HEL cells under the same treatment conditions as in (C and D). HEL-WT cells (top) exhibited morphological features consistent with early megakaryocytic differentiation including increased cytoplasmic blebbing (i), enlarged nuclei (ii), and larger cell size. In contrast, HEL-RM cells (bottom) are smaller and retain an undifferentiated, blast-like morphology characterized by a high nuclear-to-cytoplasmic ratio (iii). (**F**) HEL cells expressing RM SPOC mutants, including the individual Patch 1 alanine mutants (fuchsia), triple alanine Patch 1 mutant (Patch1Ala), and SPOC-deleted mutant (ΔSPOC), similarly retained an undifferentiated morphology. Scale bar, 50 µm.

Because mature megakaryocytes undergo endomitosis,^34,35^ we next quantified cellular polyploidization following TPA treatment of dox-inducible HEL cells. Consistent with the CD49b analyses, control cells demonstrated the expected increase in polyploidy after differentiation induction. In contrast, expression of RM substantially impaired polyploidization (**Figure 5C, S4A, and S4C**), and this defect persisted in HEL cells with the disruption of the SPOC domain. Specifically, neither the individual Patch 1 substitutions, the triple Patch 1 mutant, nor deletion of the entire SPOC domain produced a measurable restoration of high-ploidy cells (**Figure 5D**), demonstrating that SPOC disruption does not rescue this hallmark feature of megakaryocyte maturation.

To further evaluate cellular differentiation, cytospin preparations were examined for morphological features characteristic of mature megakaryocytes.^36–38^ Control HEL cells treated with TPA displayed enlarged cell size (50-100 µm), enlarged nuclei, and cytoplasmic blebbing consistent with megakaryocytic maturation (**Figure 5E and S4B**). Additionally, brightfield microscopy revealed TPA-induced cellular adhesion and spreading in control HEL cells, which were not evident in untreated cells (**Figure S4E**). In comparison, cells expressing unmutated RM retained a more immature appearance, with reduced morphological evidence of differentiation and minimal visible adhesion or spreading following TPA treatment (**Figure 5E and S4D-E**). Importantly, although cytospins from all SPOC mutant RM cell lines closely resembled those expressing unmutated RM, with no evident mature megakaryocytic morphology (**Figure 5F**), HEL-RMΔSPOC cells exhibited visible adhesion and spreading resembling TPA-treated control HEL cells (**Figure S4E**).

Together, these findings demonstrate that SPOC disruption fails to rescue RM-mediated inhibition of megakaryocytic maturation, consistent with preserved WTAP association following SPOC deletion.

### The SPOC domain contributes to RM-mediated remodeling of the m^6^A epitranscriptome

Although deletion of the SPOC domain does not abolish the interaction of RM with the MTC, the SPOC domain may still contribute to RM-dependent m^6^A deposition on specific transcripts, potentially through altered interactions with other MTC components or changes in complex stability or function. To investigate this, we performed EpiPlex m^6^A profiling (AlidaBio), a proximity barcoding-based approach that has been benchmarked against established m^6^A mapping methods and applied to m^6^A detection in independent studies.^39,40^ Poly(A)-enriched RNA from dox-induced (24 h) HEL-WT, HEL-RM, and HEL-RMΔSPOC cells was subjected to m^6^A-specific enrichment and sequencing, followed by EpiScout analysis of m^6^A profiles and integration with matched RNA-seq data (**Figure 6A and S5A-D**).

**Figure 6.**
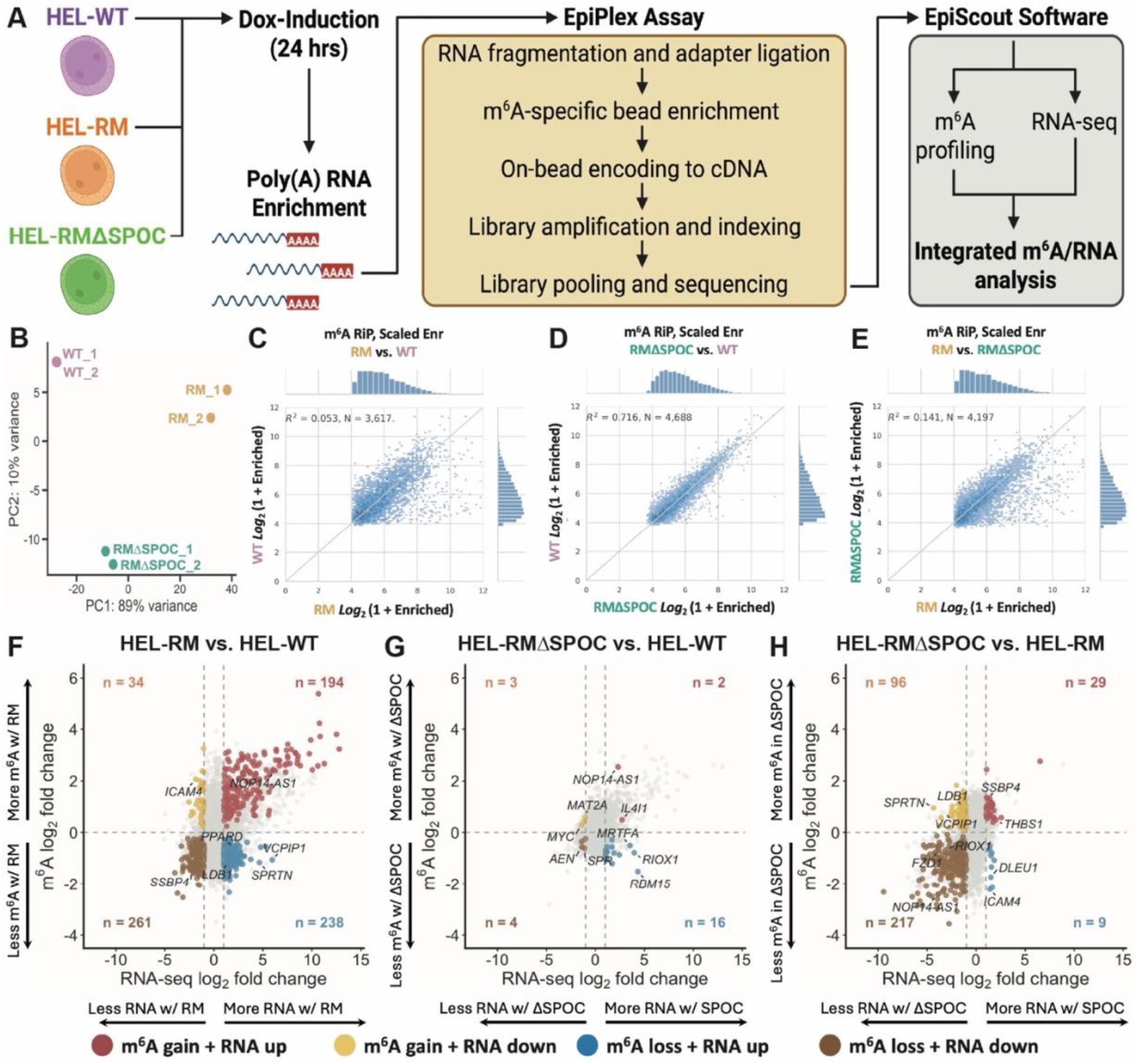
Deletion of the SPOC domain remodels the RM-associated epitranscriptomic and transcriptional landscape. (**A**) Schematic of the EpiPlex m^6^A profiling workflow. Poly(A)-enriched RNA from doxycycline-induced HEL-WT, HEL-RM, and HEL-RMΔSPOC cells was subjected to m^6^A-specific enrichment and sequencing, followed by EpiScout analysis and integration with matched RNA-seq data. Image created using BioRender.com. (**B**) Principal component (PC) analysis showing clustering of biological replicates and separation among experimental conditions (two biological replicates per condition). (**C-E**) Pairwise correlation of m^6^A enrichment between (**C**) HEL-RM versus HEL-WT, (**D**) HEL-RMΔSPOC versus HEL-WT, and (**E**) HEL-RM versus HEL-RMΔSPOC. Marginal histograms show the distribution of m^6^A enrichment values for each condition. Linear regression lines with 95% confidence intervals are shown; *R*^2^ values indicate the degree of concordance between conditions. (**F-H**) Integration of RNA expression log_2_ fold change (x-axis) and m^6^A enrichment log_2_ fold change (y-axis) for (**F**) HEL-RM versus HEL-WT, (**G**) HEL-RMΔSPOC versus HEL-WT, and (**H**) HEL-RMΔSPOC versus HEL-RM. Each point represents a gene. Genes meeting significance thresholds for both differential m^6^A enrichment (*q* < 0.05) and RNA expression effect-size threshold (|log_2_FC| ≥ 1) were classified according to the direction of m^6^A changes (e.g., gain or loss) and RNA expression changes (e.g., up or down), with the number of genes in each category indicated per quadrant. Genes with significant m^6^A changes in both directions were excluded from directional classification (see **Supplementary Table S3**). Selected genes are labeled. Vertical dashed lines denote RNA expression log_2_ fold-change thresholds of ± 1. The horizontal dashed line at an m^6^A log₂ fold change of 0 denotes no directional change in m^6^A signal and is shown as a reference rather than a statistical significance threshold.

Principal component (PC) analysis of the RNA-seq data demonstrated close clustering of biological replicates within each condition and clear separation of HEL-WT, HEL-RM, and HEL-RMΔSPOC samples (**Figure 6B**). HEL-RM separated prominently from HEL-WT along PC1, which accounted for 89% of the variance, while HEL-RMΔSPOC formed a distinct cluster much closer to HEL-WT along PC1. Pairwise comparison of m^6^A enrichment similarly showed weak correspondence between HEL-RM and HEL-WT (R^2^ = 0.053) and between HEL-RMΔSPOC and HEL-RM (R^2^ = 0.141), whereas HEL-RMΔSPOC and HEL-WT had much greater correlation (R^2^ = 0.716) (**Figure 6C-E and S5E-G**). Consistent with this global shift, among 4,174 m^6^A-enriched genomic regions matched between the RM vs. WT, and RMΔSPOC vs. WT comparisons, 3,437 (82.3%) showed a reduced absolute deviation from WT following SPOC deletion, with the median absolute m^6^A log_2_ fold-change relative to WT decreasing from 0.634 in HEL-RM to 0.255 in HEL-RMΔSPOC cells (paired Wilcoxon signed-rank test, P < 10^-300^) (**Figure S5H**). Together, these analyses demonstrate that deletion of the SPOC domain shifts the RM-associated m^6^A landscape towards a more WT-like profile.

Gene Ontology (GO) enrichment analysis revealed that genes downregulated following SPOC deletion were enriched for canonical and non-canonical WNT signaling (adjusted P = 0.0365 and 0.0228, respectively), whereas upregulated genes were enriched for hematopoietic progenitor cell differentiation (adjusted P = 0.0447; **Supplementary Table S2**). These findings suggest that SPOC deletion alters RM-associated transcriptional programs involved in signaling and hematopoietic differentiation.

We next integrated differential m^6^A enrichment (*q* < 0.05) with changes in RNA expression (|log_2_FC| ≥ 1). RM expression produced extensive changes in both m^6^A enrichment and transcript abundance relative to HEL-WT (**Figure 6F**). Among genes exhibiting increased m^6^A, 194 showed increased and 34 showed decreased RNA abundance. Conversely, among genes exhibiting m^6^A loss, 238 showed increased and 261 showed decreased RNA abundance. These findings indicate that RM-associated changes in m^6^A are not uniformly coupled to the direction of transcript abundance.

In contrast, relatively few genes had changes in both m^6^A enrichment and RNA abundance between HEL-RMΔSPOC and HEL-WT (**Figure 6G**). For HEL-RMΔSPOC, only 25 genes met both criteria, compared with 727 genes for HEL-RM.

Comparison of HEL-RMΔSPOC with HEL-RM revealed 351 genes meeting both thresholds (**Figure 6H**) with 217 having decreased m^6^A and decreased RNA abundance in HEL-RMΔSPOC compared to HEL-RM. Thus, m^6^A loss (226/351 genes) was more frequent than m^6^A gain among genes meeting both thresholds following SPOC deletion, with the largest category comprising genes with decreased m^6^A and decreased RNA abundance. Given the established role of WNT/β-catenin signaling in RM-AMKL^41^, we next examined whether genes associated with this pathway exhibited coordinated changes in m^6^A enrichment and RNA abundance following SPOC deletion. Analysis of WNT/β-catenin-associated genes revealed heterogeneous changes in m^6^A enrichment and RNA abundance across conditions, indicating that SPOC-dependent remodeling of these transcripts does not follow a uniform relationship between m^6^A and RNA expression (**Figure S5I, S6**).

Collectively, these data demonstrate that SPOC deletion substantially attenuates RM-associated m^6^A remodeling despite preserving WTAP association and failing to restore megakaryocytic differentiation, revealing a disconnect between MTC association, epitranscriptomic regulation, and the differentiation phenotype.

## Discussion

The present study challenges the prevailing model that RBM15 recruits WTAP exclusively through its SPOC domain. Although structure-guided mutagenesis demonstrated that the conserved Patch 1 basic surface mediates WTAP binding within the isolated SPOC domain, this interaction was not recapitulated in the context of the full-length RBM15. Neither the individual Patch 1 substitutions, the combined triple Patch 1 mutant, nor deletion of the entire SPOC domain abolished WTAP association with full-length RBM15 or the RM fusion protein. Together, these findings demonstrate that while the SPOC domain contributes to WTAP recognition, additional interactions maintain WTAP association with full-length RBM15 and RM. Whether these interactions involve direct protein-protein binding or are mediation by other components of the complex remains to be determined.

This discrepancy highlights the importance of studying multidomain RNA-binding proteins in their native structural context, where neighboring domains, intramolecular interactions, and conformational flexibility can influence protein function.^42^ RNA recognition motifs (RRMs), while classically associated with RNA binding, can also mediate protein-protein interactions.^28^ Combined with AlphaFold3 predictions showing spatial proximity between the RRMs and the SPOC domain, our data raise the possibility that one or more RRMs contribute directly or indirectly to WTAP recruitment. Consistent with distinct functions of RBM15 domains in RNA regulation, Yu et al. recently demonstrated that RBM15 promotes circular RNA biogenesis through cooperative activities of its SPOC domain and intrinsically disordered domains, whereas RRM2 and RRM3 contribute to nuclear retention of circular RNAs. Notably, the effect of circular RNA biogenesis was independent of METTL3, further supporting m^6^A-independent functions of RBM15.^43^ Identifying these determinants will be important for defining the molecular architecture underlying RBM15-MTC assembly.

Importantly, however, the preservation of MTC association following SPOC deletion did not equate to preservation of the RM-associated epitranscriptome. EpiPlex profiling revealed extensive differences in m^6^A enrichment between HEL-RM and HEL-WT, whereas deletion of SPOC shifted the m^6^A landscape toward a more WT-like state. Direct comparison of matched m^6^A regions relative to WT further demonstrated that 82.3% exhibited reduced deviation from WT following SPOC deletion, indicating that loss of SPOC broadly attenuates the magnitude of RM-associated m^6^A alterations. Integration with RNA-seq nevertheless revealed heterogeneous relationships between changes in m^6^A enrichment and transcript abundance, indicating that SPOC-dependent epitranscriptomic remodeling is not uniformly coupled to a direction of gene-expression changes. These findings suggest that the SPOC domain may be dispensable for biochemical association with the MTC but nevertheless important for MTC activity or targeting at specific transcripts. Thus, physical association with WTAP alone may not be sufficient to reproduce the RM-associated m^6^A landscape.

SPOC deletion remodeled m^6^A and transcription without restoring megakaryocytic differentiation. This apparent disconnect suggests that the differentiation block imposed by RM may not require preservation of the entire RM-associated epitranscriptomic program. Residual methylation of critical targets, SPOC-independent functions of RM, or parallel transcriptional mechanisms may therefore be sufficient to maintain the immature phenotype. These findings caution against equating molecular rescue with phenotypic rescue and suggest that defining the specific m^6^A-regulated transcripts required for RM leukemogenic activity will be an important next step.

Several questions remain. Although our findings establish SPOC-independent WTAP association, the responsible interaction surfaces remain undefined and require systematic structural and biochemical investigation. Furthermore, while the integration of EpiPlex and RNA-seq identifies associations between SPOC-dependent changes in m^6^A enrichment and transcript abundance, these analyses do not establish a direct causal relationship between methylation and expression of individual transcripts. Orthogonal validation of candidate m^6^A targets and functional interrogation of their contribution to RM-mediated leukemogenesis will therefore be important.

Collectively, our findings demonstrate that WTAP association, SPOC-dependent m^6^A remodeling, and RM-mediated inhibition of megakaryocytic differentiation are functionally separable. Although the SPOC domain is dispensable for stable WTAP association, its deletion substantially attenuates RM-associated changes in the m^6^A epitranscriptome without rescuing the differentiation block in HEL cells. These findings challenge a model in which SPOC-dependent WTAP recruitment is the principal determinant of RM-mediated RNA regulation and suggest that the SPOC domain contributes to the functional organization or targeting of the m^6^A machinery. Importantly, the persistence of impaired megakaryocytic differentiation despite extensive restoration of a wild-type-like m^6^A landscape suggests that additional or more selective molecular mechanisms sustain the RM-associated differentiation block.

## Acknowledgements

The authors thank Ananya Anmangandla, Michael Chabot, and Jerome Santos of AlidaBio for their expertise and guidance regarding the EpiPlex assay and EpiScout software. They thank Maximillian J. Carlino for providing the nucleofection buffer used to generate the mutant HEL-RM cell lines. They thank the Yale Flow Cytometry Core (supported, in part, by a National Institutes of Health [NIH] Cancer Center Support Grant [NIH P30CA016359]) and the Yale Center for Genome Analysis (supported by the NIH, National Institute of General Medical Sciences [NIGMS] under award number 1S10OD030363-01A1) for assistance with sequencing. Molecular graphics and analyses of RBM15 protein structure were performed using the University of California, San Francisco (UCSF) ChimeraX software, developed by the Resource for Biocomputing, Visualization, and Informatics at UCSF with support from the NIH NIGMS grant P41GM103311.

J.A.E. received funding from the American Society of Hematology (ASH) Hematology Inclusion Pathway Graduate Student Award, the Lo Graduate Fellowship for Excellence in Stem Cell Research, the NIH National Heart, Lung, and Blood Institute (NHLBI) grant T32HL007974, and the NIH National Cancer Institute (NCI) grant F31CA309946. M.Y.M. received funding from the Yale Cooperative Center of Excellence in Hematology (YCCEH) NIH National Institute of Diabetes and Digestive and Kidney Diseases (NIDDK) grant U54DK106857, the Lo Graduate Fellowship for Excellence in Stem Cell Research, and the NIH NCI grant F31CA271571. H.M. received funding from YCCEH NIH NIDDK grant U54DK106857. N.A.-L. received funding from the NIH NHLBI grant T32HL007974. D.S.K. received funding from the Alex’s Lemonade Stand Foundation (ALSF) Innovation Grant 1433524, the NIH NCI grant R01CA222518, the NIH NIDDK grant RC2DK122376, and the YCCEH NIH NIDDK grant U54DK106857.

## Authorship Contributions

J.A.E. and D.S.K. conceptualized the study; J.A.E., M.Y.M., and M.C. were responsible for data curation; J.A.E. conducted formal analysis; J.A.E., M.Y.M., and D.S.K. acquired funding; J.A.E., M.Y.M., M.C., H.M., G.L., L.W., P.Z., and N.A.-L. conducted the investigation; J.A.E., M.Y.M., and D.S.K. designed the methodology; J.A.E. and D.S.K. were responsible for project administration; D.S.K. provided resources; J.A.E. and D.S.K. provided supervision; J.A.E., M.Y.M., M.C., H.M., and G.L. validated the results; J.A.E. was responsible for visualizing the data; J.A.E. and D.S.K. wrote the original draft; and J.A.E., M.Y.M., M.C., H.M., G.L., L.W., P.Z., N.A.-L., and D.S.K. reviewed and edited the manuscript.

## Conflict of Interest Disclosures

The authors declare no competing financial interests.

## Notes

### Competing Interest Statement

The authors have declared no competing interest.

