## Supplementary Figures and Table for "RBM15::MKL1 uncouples SPOC-dependent m^6^A remodeling from WTAP recruitment"

**Short Title:** SPOC-dependent m$^6$A remodeling by RBM15::MKL1

**Authors:**

Jether Amos Espinosa[1,2], Mi Chen[1,3], Madeline Y. Mayday[1,2], Hummi Mohammed[1], Gina Lee[1,4], Lin Wang[1,3], Ping-Xia Zhang[1,3], Nadia Ayala-Lopez[1,3,§], and Diane S. Krause[1,2,3,5,6*]

**Affiliations:**

[1]Yale Stem Cell Center, [2]Department of Pathology, [3]Department of Laboratory Medicine, [4]Department of Molecular, Cellular, and Developmental Biology, [5]Department of Cell Biology and [6]Yale Cancer Center, New Haven, Connecticut, USA.

[§]Present address: Labcorp Bioanalytical Services, Indianapolis, Indiana, USA.

**Data Sharing Statement:**

All data generated in this study are included in this article and its supplementary information. RNA-seq and EpiPlex data have been deposited in the Gene Expression Omnibus (GEO) repository and are accessible with the accession number GSE349759.

**Supplementary Information:**

| Immunoblotting: | | | | |
|---|---|---|---|---|
| **Target** | **Host Species** | **Dilution Factor** | **Vendor** | **Catalog Number** |
| FLAG | Mouse | 1:2000 | Millipore Sigma | F1804-200UG |
| Lamin B1 | Mouse | 1:5000 | Proteintech | 66095-1-1g |
| METTL14 | Rabbit | 1:1000 | Proteintech | 26158-1-AP |
| METTL3 | Rabbit | 1:1000 | Bethyl Laboratories | A301-567A |
| Mouse species | Mouse | 1:2000 | Santa Cruz Biotechnology | SC-516102 |
| Rabbit species | Mouse | 1:2000 | Santa Cruz Biotechnology | SC-2357 |
| Rat | Goat | 1:2000 | Invitrogen | 62-9520 |
| WTAP | Rabbit | 1:1000 | Abcam | AB195380 |
| ZC3H13 | Rabbit | 1:2000 | Proteintech | 28430-1-AP |
| **Immunoprecipitation:** | | | | |
| **Target** | **Host Species** | **Dilution Factor** | **Vendor** | **Catalog Number** |
| FLAG | Rat | 1:22.5 | BioLegend | 637303 |
| **Flow Cytometry:** | | | | |
| **Target** | **Target Species** | **Dilution Factor** | **Vendor** | **Catalog Number** |
| APC-CD49b | Human | 1:100 | BioLegend | 359310 |
| PE-CD49b | Human | 1:100 | BioLegend | 359307 |

**Table S1: Antibodies used in this study.**

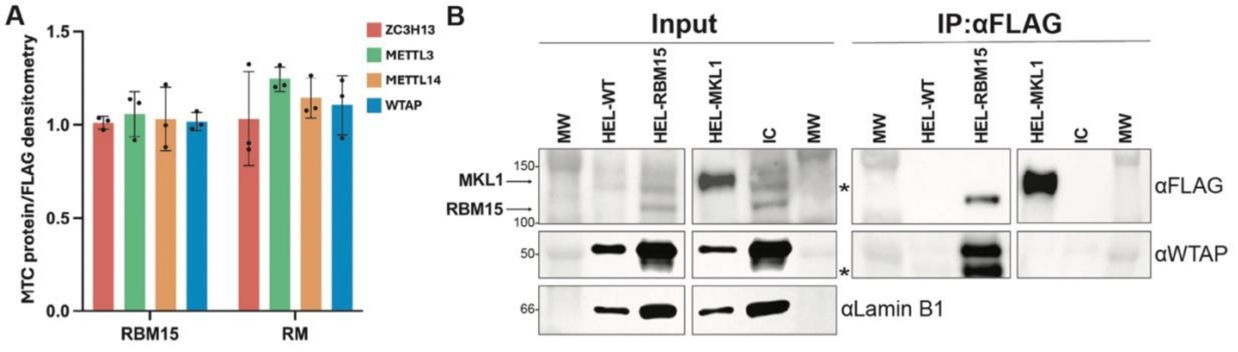


**Figure S1. Methyltransferase complex (MTC) association with RBM15 and RM, related to Figure 1.** (**A**) Densitometric quantification of ZC3H13, METTL3, METTL14, and WTAP co-immunoprecipitated with FLAG-tagged RBM15 or RBM15::MKL1 (RM).

MTC protein signals were normalized to their corresponding FLAG pulldown and subsequently normalized to the RBM15_1 control within each independent experiment. Technical duplicate RBM15 and RM samples were averaged within each experiment. Bars represent mean ± standard error of the mean (SEM) from three independent experiments; individual experiments are shown as points. (**B**) Anti-FLAG immunoprecipitation (IP) of nuclear lysates from dox-inducible HEL-WT, HEL-RBM15, and HEL-MKL1 cells. Input and FLAG-IP fractions were immunoblotted for FLAG and endogenous WTAP. FLAG-tagged RBM15 co-immunoprecipitated WTAP, whereas FLAG-tagged MKL1 did not detectably associate with WTAP, indicating that WTAP association is not conferred by MKL1. Lamin B1 served as a nuclear input control. Asterisks (*) indicate nonspecific bands. IgG isotype control (IC) for the pulldown is from nuclear lysate of dox-induced HEL-RBM15. Molecular weight (MW) markers are indicated in kDa.

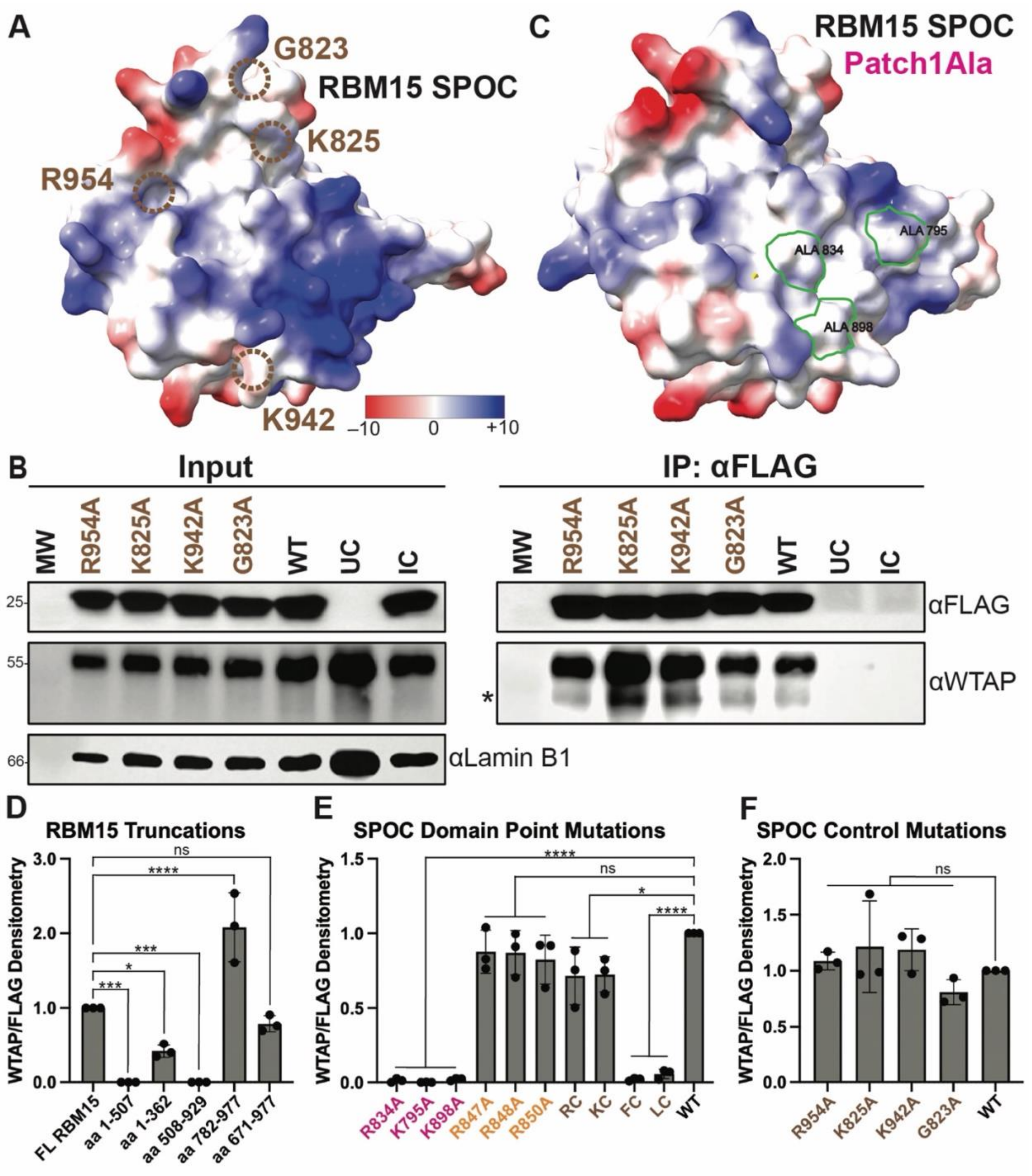


**Figure S2. Additional information on RBM15 SPOC domain mutations, related to Figure 2.** (**A**) AlphaFold3 (AF3) structural model of the RBM15 SPOC domain. Electrostatic surface potential was visualized using the Coulombic Surface Coloring tool in UCSF ChimeraX[22] and is depicted ranging from –10 (red), 0 (white), to +10 (blue) kcal/(mol*e). The approximate locations of the four additional residues outside Patch 1 are noted by dashed brown circles. The structure represents the unmutated RBM15 SPOC domain. (**B**) Immunoblot showing that the additional alanine mutants referenced in (A) do not abrogate WTAP co-immunoprecipitation (co-IP), similar to unmutated SPOC (WT). Untransfected HEK293T cells were included as a control (UC). Nuclear fraction of cell lysates was used for co-IP and, with Lamin B1 serving as a nuclear loading control for the input. IgG isotype control (IC) for the pulldown is from nuclear lysate of HEK293T transfected with the WT construct. Molecular weight (MW) markers are indicated in kDa. Representative blot is shown (n = 3 biological replicates). Asterisk (*) indicates non-specific bands. (**C**) AF3 structural model of the triple Patch 1 mutant (R834A, K795A, and K898A) in the RBM15 SPOC domain visualized as in (A). Loss of Patch 1 electrostatic potential is outlined in green. (**D**-**F**) Densitometric quantifications of WTAP co-immunoprecipitation normalized to the corresponding FLAG pulldown for (**D**) RBM15 truncation mutants related to figure 2B, (**E**) individual SPOC domain alanine mutants related to figure 2E, and (**F**) additional alanine mutants related to supplementary figure 2B. Data are presented as mean ± standard error of the mean (SEM). Values in (D) are expressed relative to full-length (FL) RBM15 (FL RBM15 = 1.0), whereas values in (E) and (F) are expressed relative to WT SPOC (WT = 1.0).

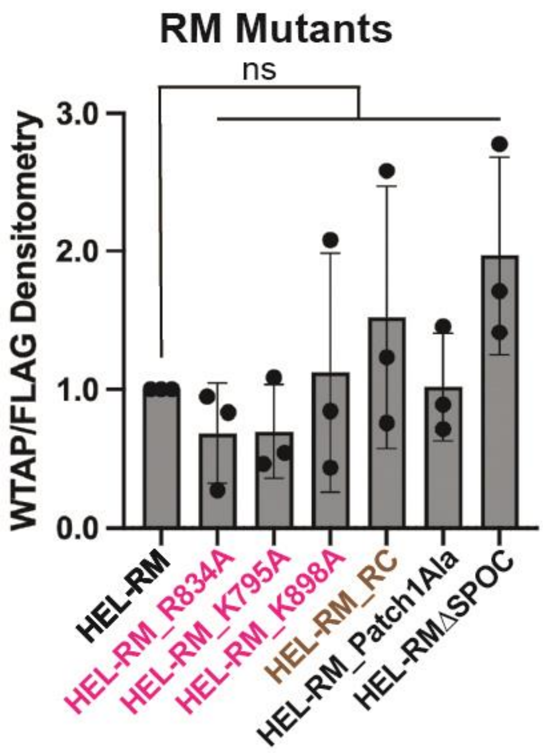


**Figure S3. Additional information on RM mutants, related to Figure 4.** Densitometric quantification of WTAP co-immunoprecipitation normalized to the corresponding FLAG pulldown for RM containing individual alanine substitutions (R834A, K795A, K898A, and R886A non-Patch 1 control [RC]), the triple Patch 1 alanine mutant (Patch1Ala), and the SPOC domain-deletion mutant (ΔSPOC). Data are presented as mean ± standard error of the mean (SEM) from three biological replicates. Values are expressed relative to HEL-RM (HEL-RM = 1.0).

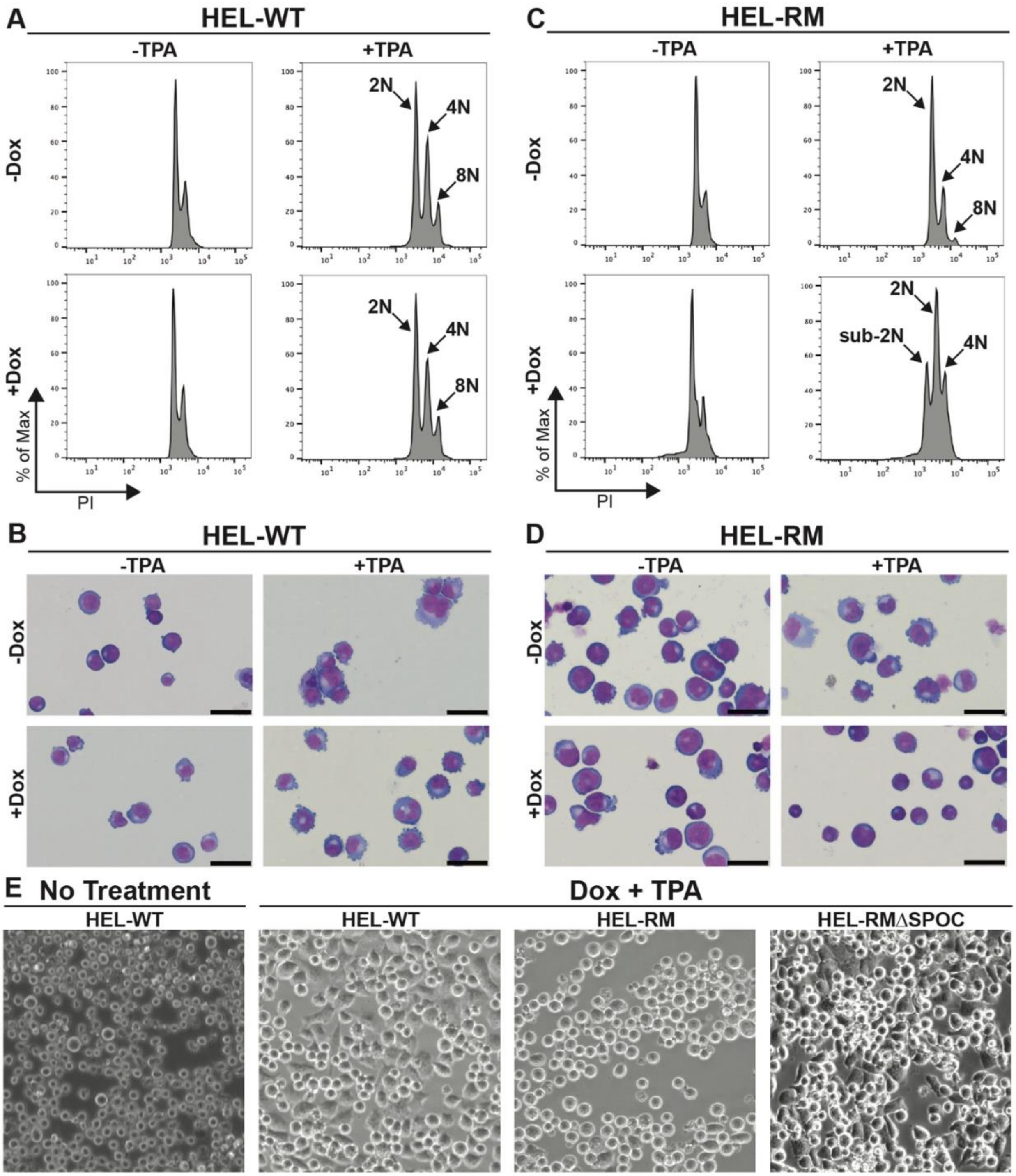


**Figure S4. Additional information on HEL cell differentiation, related to Figure 5.** (**A**) Flow cytometry histograms showing polyploidization peaks of propidium iodide (PI)-stained HEL-WT cells treated with or without doxycycline (dox) or 12-*O*-tetradecanoylphorbol-13-acetate (TPA) for 72 hours. Ploidy peaks of 2N, 4N, and 8N are noted. (**B**) Representative Wright-Giemsa-stained cytospins of HEL-WT cells treated with the same conditions as (A). Dox treatment shows no effect in blocking TPA-induced differentiation in HEL-WT cells. (**C**) Flow cytometry histograms showing polyploidization peaks of PI-stained HEL-RM cells treated with the same conditions as (A). Ploidy peaks of 2N, 4N, and 8N are noted in TPA-induced cells, indicating differentiation (upper-right panel). The presence of dox-induced RM expression blocks TPA-induced differentiation, as indicated by the presence of a sub 2N peak and absence of an 8N peak (bottom-right panel). (**D**) Representative Wright-Giemsa-stained cytospins of HEL-RM cells treated with the same conditions as (A). Dox treatment alone does not alter the differentiation phenotype (bottom-left panel), but the presence of dox-induced RM blocks TPA-induced differentiation (bottom-right panel). Scale bar, 50 μm. (**E**) Representative phase-contrast images of untreated HEL-WT cells and HEL-WT, HEL-RM, and HEL-RMΔSPOC cells treated with dox and TPA for 24 hours. Untreated HEL-WT cells exhibited predominantly rounded, nonadherent morphology, whereas dox- and TPA-treated HEL-WT cells displayed visible cell adhesion and spreading. In contrast, dox- and TPA-treated HEL-RM cells retained a predominantly rounded, nonadherent morphology resembling intreated HEL-WT cells. Notably, HEL-RMΔSPOC cells exhibited morphological features resembling treated HEL-WT cells, including visible adhesion and spreading.

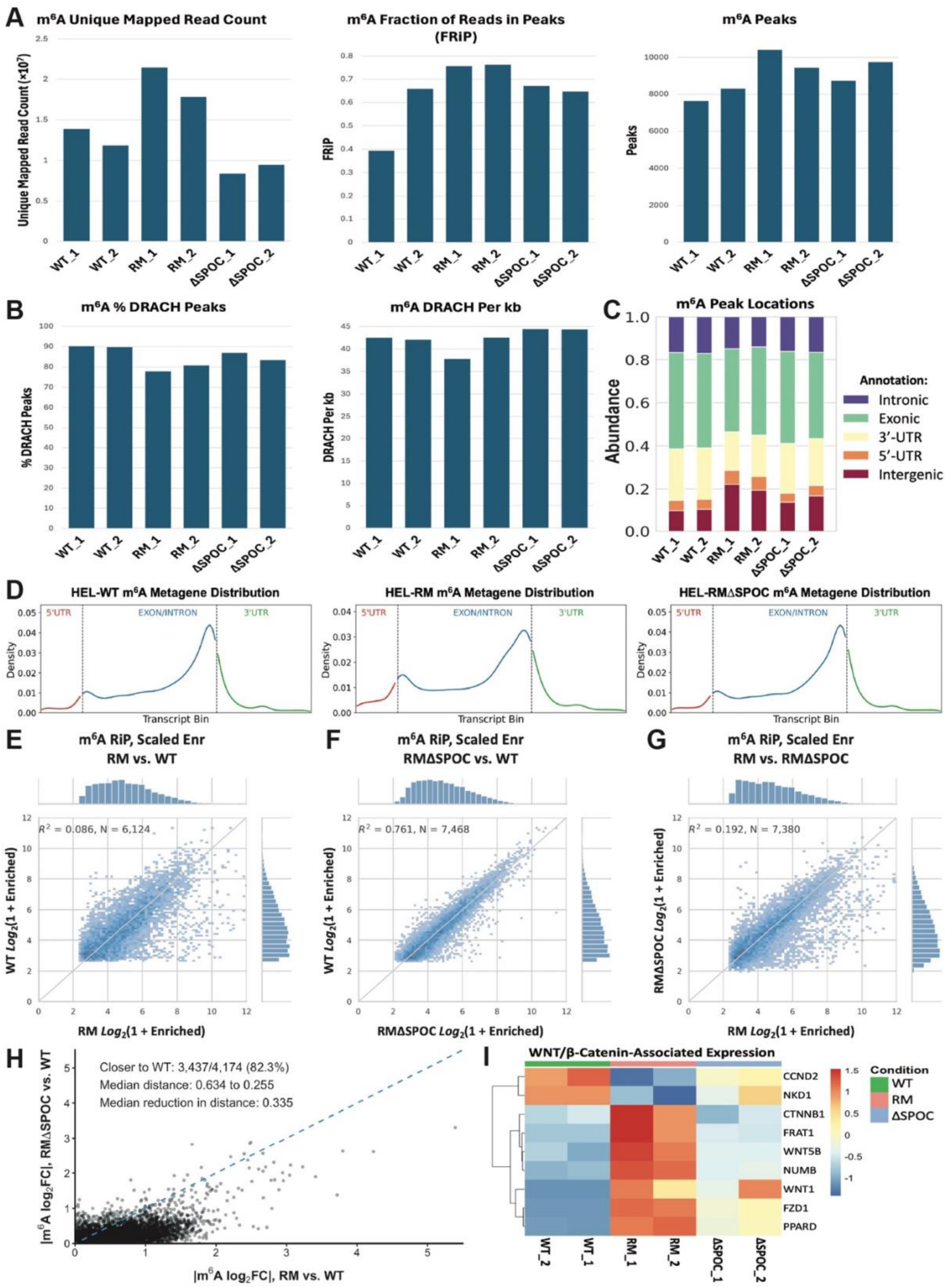
A
m6A Unique Mapped Read Count
Unique Mapped Read Count (×10⁷)
m6A Fraction of Reads in Peaks (FRiP)
FRiP
m6A Peaks
Peaks
WT_1
WT_2
RM_1
RM_2
ΔSPOC_1
ΔSPOC_2
B
m6A % DRACH Peaks
% DRACH Peaks
m6A DRACH Per kb
DRACH Per kb
C
m6A Peak Locations
Abundance
Annotation:
Intronic
Exonic
3'-UTR
5'-UTR
Intergenic
D
HEL-WT m6A Metagene Distribution
HEL-RM m6A Metagene Distribution
HEL-RMΔSPOC m6A Metagene Distribution
5'UTR
EXON/INTRON
3'UTR
Density
Transcript Bin
E
m6A RiP, Scaled Enr
RM vs. WT
R² = 0.086, N = 6,124
WT Log2(1 + Enriched)
RM Log2(1 + Enriched)
F
m6A RiP, Scaled Enr
RMΔSPOC vs. WT
R² = 0.761, N = 7,468
WT Log2(1 + Enriched)
RMΔSPOC Log2(1 + Enriched)
G
m6A RiP, Scaled Enr
RM vs. RMΔSPOC
R² = 0.192, N = 7,380
RMΔSPOC Log2(1 + Enriched)
RM Log2(1 + Enriched)
H
Closer to WT: 3,437/4,174 (82.3%)
Median distance: 0.634 to 0.255
Median reduction in distance: 0.335
|m6A log2FC|, RMΔSPOC vs. WT
|m6A log2FC|, RM vs. WT
I
WNT/β-Catenin-Associated Expression
CCND2
NKD1
CTNNB1
FRAT1
WNT5B
NUMB
WNT1
FZD1
PPARD
Condition
WT
RM
ΔSPOC
WT_2
WT_1
RM_1
RM_2
ΔSPOC_1
ΔSPOC_2

**Figure S5. Quality control and global characterization of EpiPlex m$^6$A and RNA-seq datasets, related to Figure 6.** (**A**) Quality control metrics from the full-depth EpiPlex m$^6$A-enriched datasets for two biological replicates each of wild-type (WT) HEL cells, those expressing RBM15::MKL1 (RM), or RM lacking the SPOC domain (ΔSPOC). Bar graphs show the number of uniquely mapped m$^6$A reads, fraction of reads in peaks (FRiP), and the number of identified m$^6$A peaks for each sample. (**B**) Sequence-motif characteristics of m$^6$A peaks from the full-depth EpiPlex datasets, showing the percentage of peaks containing the consensus DRACH motif and the density of DRACH motifs per kilobase (kb). (**C**) Genomic distribution of m$^6$A peaks from the full-depth EpiPlex datasets, categorized as intronic, exonic, 3' untranslated region (3'-UTR), 5' untranslated region (5'-UTR), or intergenic. (**D**) Metagene distributions of m$^6$A enrichment across transcripts in WT, RM, and ΔSPOC cells generated using the full-depth datasets. m$^6$A peak distributions are mapped onto a hypothetical transcript normalized to 100 bins, illustrating enrichment across the 5'-UTR, exon/intron-containing transcript body, and 3'-UTR. Experimentally determined 5'-UTR and 3'-UTR boundaries are indicated at binds 10 and 60, respectively. (**E-G**) Pairwise correlations of scaled m$^6$A enrichment from full-depth EpiPlex datasets comparing (**E**) RM versus WT, (**F**) ΔSPOC versus WT, and (**G**) RM versus ΔSPOC. Marginal histograms show the distribution of m$^6$A enrichment values for each condition. Linear regression lines with 95% confidence intervals are shown; $R^2$ values indicate the degree of concordance between conditions. (**H**) Comparison of the absolute magnitude of m$^6$A enrichment changes relative to WT across 4,174 genomic regions matched between the HEL-RM versus HEL-WT and HEL-RMΔSPOC versus HEL-WT datasets. Each point represents one matched m$^6$A region. The blue dashed identity line denotes equal absolute deviation from WT in HEL-RM and HEL-RMΔSPOC; points below the line indicate regions that are closer to WT following SPOC deletion. Of the 4,174 matched regions, 3,437 (82.3%) showed reduced deviation from WT in HEL-RMΔSPOC cells. The median absolute distance from WT decreased from 0.634 in HEL-RM to 0.255 in HEL-RMΔSPOC, with a median reduction in distance of 0.335 $\log_2$FC. (**I**) Heatmap of RNA-seq expression data for selected WNT/β-catenin-associated genes across WT, RM, and ΔSPOC biological replicates. Expression values are shown as a row-scaled relative abundance. Two biological replicates were analyzed per condition.

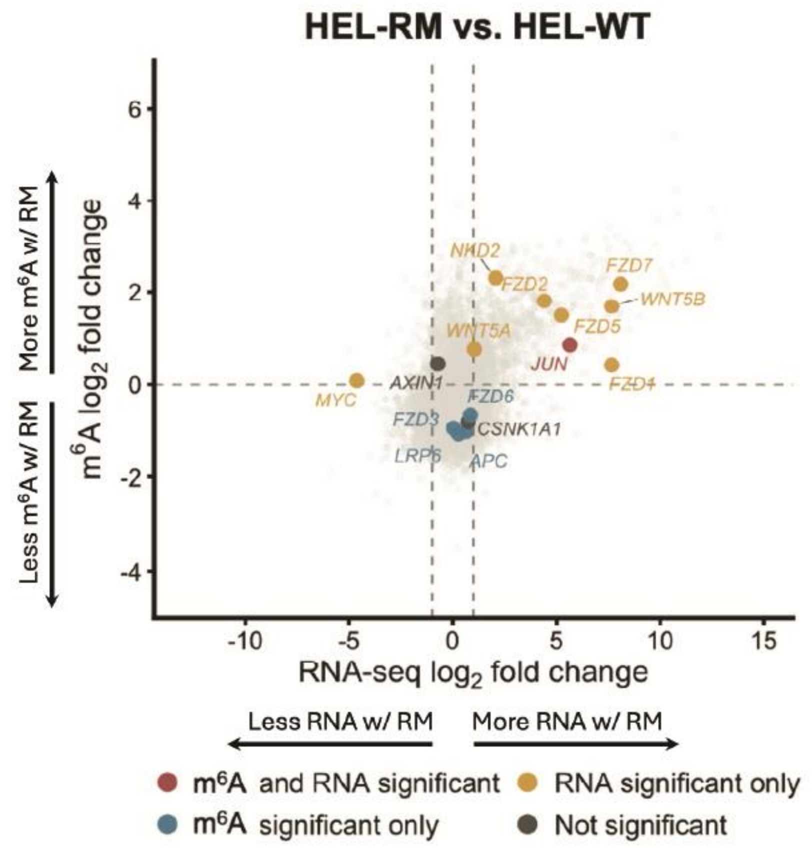


**Figure S6. Integrated m$^6$A and RNA-seq analysis of WNT/β-catenin pathway-associated genes, related to Figure 6F.** Integrated analysis of differential m$^6$A modification and gene expression in doxycycline-induced HEL cells expressing RBM15::MKL1 (HEL-RM) relative to wild-type control cells (HEL-WT). Each point represents a gene detected in both the m$^6$A and RNA-seq datasets, with RNA-seq $\log_2$ fold change (HEL-RM vs. HEL-WT) plotted on the x-axis and m$^6$A $\log_2$ fold change plotted on the y-axis. WNT/β-catenin pathway-associated genes are highlighted and labeled, whereas all other genes in the integrated dataset are shown in light gray for reference. Highlighted genes are categorized according to statistical significance in the two datasets: red, significant differential m$^6$A modification and RNA expression; yellow, significant RNA expression only; blue, significant differential m$^6$A modification only; and dark gray, not significant in either analysis. RNA-seq significance was defined as an adjusted $P$ value $< 0.05$ and an absolute $\log_2$ fold change $\geq 1$, whereas $m^6A$ significance was defined as a $q$ value $< 0.05$. Vertical dashed lines denote RNA-seq $\log_2$ fold changes of $-1$ and $+1$, corresponding to the twofold expression-change threshold. The horizontal dashed line at an $m^6A$ $\log_2$ fold change of 0 denotes no directional change in $m^6A$ signal and is shown as a reference rather than a statistical significance threshold. Positive values indicate increased RNA expression or $m^6A$ signal in HEL-RM relative to HEL-WT, whereas negative values indicate decreased RNA expression or $m^6A$ signal. WNT/β-catenin pathway-associated genes were selected to include WNT ligands and receptors, core pathway regulators, and downstream pathway-associated genes.
